# Metabolomic consequences of genetic inhibition of PCSK9 compared with statin treatment

**DOI:** 10.1101/278861

**Authors:** Eeva Sliz, Johannes Kettunen, Michael V Holmes, Clare Oliver-Williams, Charles Boachie, Qin Wang, Minna Männikkö, Sylvain Sebert, Robin Walters, Kuang Lin, Iona Y Millwood, Robert Clarke, Liming Li, Naomi Rankin, Paul Welsh, Christian Delles, J. Wouter Jukema, Stella Trompet, Ian Ford, Markus Perola, Veikko Salomaa, Marjo-Riitta Järvelin, Zhengming Chen, Debbie A Lawlor, Mika Ala-Korpela, John Danesh, George Davey Smith, Naveed Sattar, Adam Butterworth, Peter Würtz

## Abstract

**Background:** Both statins and PCSK9 inhibitors lower blood low-density lipoprotein cholesterol (LDL-C) levels to reduce risk of cardiovascular events. To assess potential differences between metabolic effects of these two lipid-lowering therapies, we performed detailed lipid and metabolite profiling of a large randomized statin trial, and compared the results with the effects of genetic inhibition of PCSK9, acting as a naturally occurring trial.

**Methods:** 228 circulating metabolic measures were quantified by nuclear magnetic resonance spectroscopy, including lipoprotein subclass concentrations and their lipid composition, fatty acids, and amino acids, for 5,359 individuals (2,659 on treatment) in the PROspective Study of Pravastatin in the Elderly at Risk (PROSPER) trial at 6-months post-randomization. The corresponding metabolic measures were analyzed in eight population cohorts (N=72,185) using *PCSK9* rs11591147 as an unconfounded proxy to mimic the therapeutic effects of PCSK9 inhibitors.

**Results:** Scaled to an equivalent lowering of LDL-C, the effects of genetic inhibition of PCSK9 on 228 metabolic markers were generally consistent with those of statin therapy (*R*^2^=0.88). Alterations in lipoprotein lipid composition and fatty acid balance were similar. However, discrepancies were observed for very-low-density lipoprotein (VLDL) lipid measures. For instance, genetic inhibition of PCSK9 showed weaker effects on lowering of VLDL-cholesterol compared with statin therapy (54% vs. 77% reduction, relative to the lowering effect on LDL-C; *P*=2 × 10^−7^ for heterogeneity). Genetic inhibition of PCSK9 showed no robust effects on amino acids, ketones, and a marker of inflammation (GlycA); in contrast, statin treatment lowered GlycA levels.

**Conclusions:** Genetic inhibition of PCSK9 results in similar metabolic effects as statin therapy across a detailed lipid and metabolite profile. However, for the same lowering of LDL-C, PCSK9 inhibitors are predicted to be less efficacious than statins at lowering VLDL lipids, which could potentially translate into subtle differences in cardiovascular risk reduction.

## INTRODUCTION

Statins are first line therapy to lower blood levels of low-density lipoprotein cholesterol (LDL-C) and reduce the risk of cardiovascular events (1–3). Treatment with proprotein convertase subtilisin/kexin type 9 (PCSK9) inhibitors has emerged as an additional effective therapy to lower LDL-C, resulting in reductions of approximately 45–60% (4, 5). Large cardiovascular outcome trials have recently demonstrated that PCSK9 inhibitors reduce the risk of major cardiovascular events when added to statin treatment (6, 7). Based on the first major outcome trial (6), there has been some suggestions that PCSK9 inhibitors may be slightly less efficacious as statins for a given LDL-C reduction; however, other reports suggest that this is not the case, with apparent differences in cardiovascular event reduction explained by the short duration of the PCSK9 trials (8). Assessment of the detailed metabolic effects of statins and PCSK9 inhibitors could provide a more detailed understanding of these lipid-lowering therapies and may shed light on potential discrepancies in their effects on lipid and lipoprotein metabolism.

The anticipated pharmacological effects of PCSK9 inhibitors may be assessed by LDL-C lowering variants in the *PCSK9* gene, which act as unconfounded proxies for the lifetime effects of treatments (9–11). The observation of a prominent lower risk of coronary heart disease for LDL-C lowering variants in *PCSK9* was pivotal for accelerating the development of anti-PCSK9 therapeutics (10).supporting the validity of using genetic proxies for molecular characterization of lipid-lowering targets, we have previously shown that LDL-C lowering variants in *HMGCR* (the gene encoding the target for statins) closely recapitulate the fine-grained metabolic changes associated with starting statin therapy, as assessed by nuclear magnetic resonance (NMR) metabolomics in longitudinal cohorts (12). These intricate metabolic effects of statins were recently confirmed in PREVEND IT (Prevention of Renal and Vascular End-stage Disease Intervention Trial), a small randomized trial (13). Other studies have assessed the associations of *PCSK9* variants with lipoprotein subclass profiles (14, 15), and the treatment effects of PCSK9 inhibitors on lipoprotein particle concentrations and lipidomic measures have been examined in small trials (16–18). However, prior studies have had limited power to dissect potential differences between PCSK9 inhibition and statin therapy for the same LDL-C lowering conferred, complicating direct comparisons of their impact on detailed lipid and metabolite measures.

In the present study, we examined the effects of statin therapy and genetic inhibition of PCSK9 on a circulating metabolic profile of 228 metabolic measures, quantified by NMR metabolomics, including lipoprotein subclasses, their lipid concentrations and composition, fatty acid balance, and several non-lipid pathways. The metabolic effects of statin treatment were assessed in a large randomized controlled trial. In the absence of NMR metabolomics data from a large randomized trial of PCSK9 inhibitor therapy, the anticipated pharmacological effects were examined for a loss-of-function variant in the *PCSK9* gene (10, 19). Comparing the metabolomic effects of genetic inhibition of PCSK9 to statin therapy provides an opportunity to examine possible discrepancies in many circulating biomarkers, and in turn elucidate potential therapeutic differences in the molecular mechanisms to reduce cardiovascular risk.

## METHODS

### Study design

An overview of the study design is shown in **Figure 1**. NMR metabolomics was performed on 5,359 blood samples from the PROSPER (PROspective Study of Pravastatin in the Elderly at Risk) trial (20) at 6-month post-randomization, and 72,185 samples from eight population cohorts from the United Kingdom (INTERVAL (21), Avon Longitudinal Study of Parents (ALSPAC) mothers and offspring (22, 23)), Finland (FINRISK-1997, FINRISK-2007 (24), and Northern Finland Birth Cohort studies 1966 and 1986 (25, 26)), and China (China Kadoorie Biobank (27)). All study participants provided written informed consent, and study protocols were approved by the local ethics committees.

**Figure 1.**
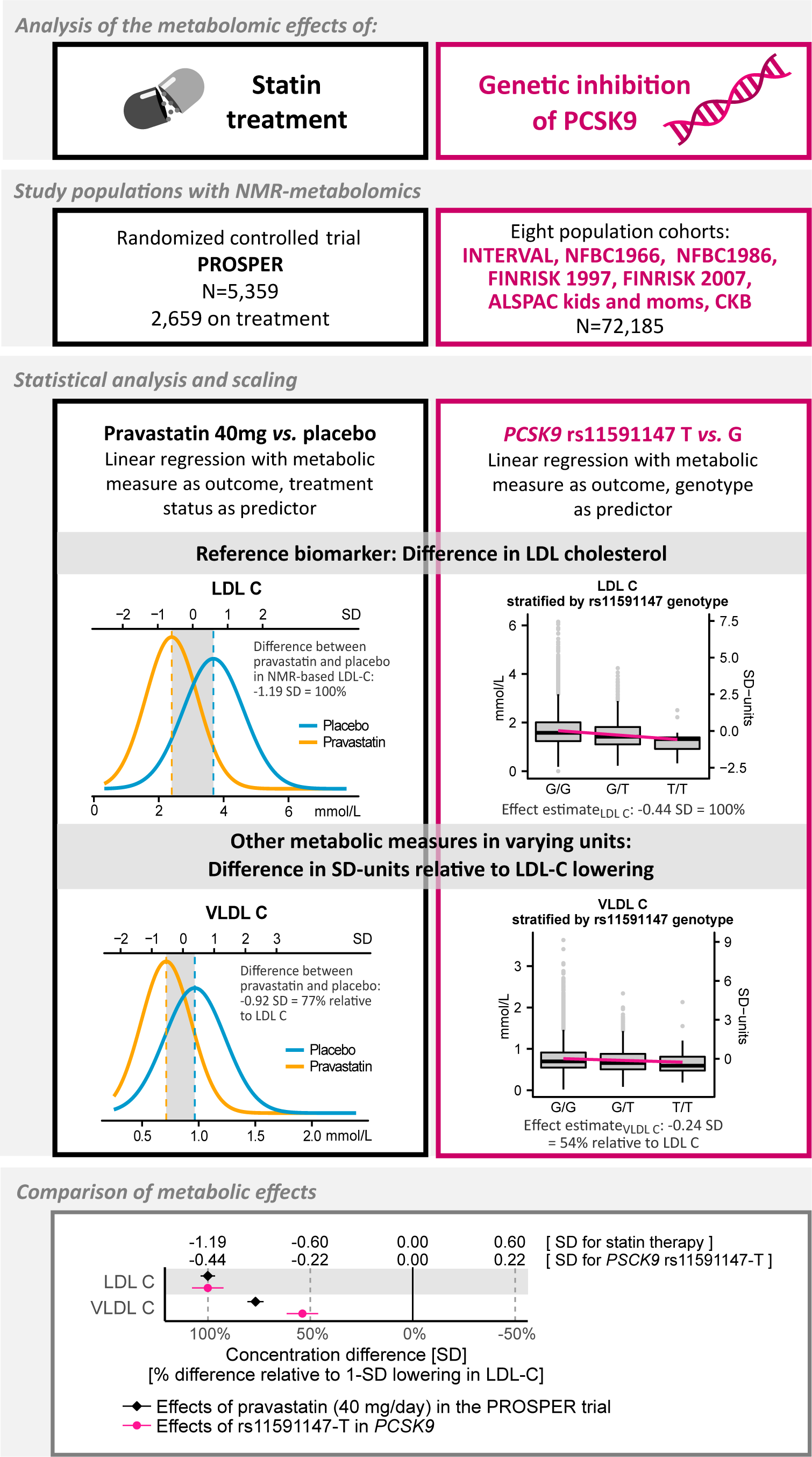
Overview of the study design and statistical analyses.

PROSPER is a double-blind, randomized placebo controlled trial investigating the benefit of pravastatin (40 mg/day) in elderly individuals at risk of cardiovascular disease, with 5,804 participants (70–82 years old) from Scotland, Ireland and the Netherlands enrolled between December 1997 and May 1999 (28). All participants had above average plasma total cholesterol concentration (4.0 to 9.0 mmol/L) at baseline and 50% had prior vascular disease. For the present study, 5,359 samples (2,659 on pravastatin) were measured by NMR metabolomics; all were previously unthawed 6-month post-randomization EDTA plasma samples stored at −80°C (28). Metabolite data from baseline samples were not available, however the randomization should ensure that there are limited between-group differences at baseline. Replication of the metabolic effects of pravastatin in PROSPER was done by comparison with recent results from PREVEND-IT (13).

The metabolic effects of PCSK9 inhibition were assessed via the principle of Mendelian randomization using rs11591147-T (R46L), a loss-of-function allele robustly associated with lower LDL-C and decreased cardiovascular risk (10, 11). Additional genetic variants in the *PCSK9* locus, which have previously been used in Mendelian randomization studies on *PCSK9* (11, 29) and display low linkage disequilibrium with rs11591147 (*R*^2^<0.2) were assessed in sensitivity analyses. To complement the comparison of *PCSK9* rs11591147 effects against the statin trial, we further examined the metabolic effects of rs12916 in *HMGCR* in the same study population (N=72,185), acting as a trial of a very small statin dose by naturally occurring randomization of HMG-CoA reductase inhibition (12, 30). Among SNPs in *HMGCR*, rs12916 exhibits the strongest association with LDL-C and has been shown to affect hepatic HMGCR expression as well as cardiovascular risk (11, 12, 30). Finally, to corroborate the validity of using genetic proxies to mimic the randomized trial effects, we compared metabolic effects of statin treatment in PROSPER with the corresponding effects of *HMGCR* rs12916. Pregnant women and individuals on lipid-lowering treatment were excluded from the analyses where information was available. Details of the cohorts are provided in **Supplementary Methods** and **Supplementary Table 1**.

**Table 1.** Baseline characteristics of participants in the PROSPER statin trial and cohorts for analyses of genetic inhibition of PCSK9.

| Characteristics | PROSPER statin trial |  | Cohorts in analyses of genetic variants |
| --- | --- | --- | --- |
|  | Placebo | Pravastatin |  |
| Number of individuals | 2,700 | 2,659 | 72,185 |
| Male (%) | 48.3 | 48.3 | 45.2 |
| Age (year) | 75.3 ± 3.4 | 75.4 ± 3.3 | 33.1 ± 6.7 |
| BMI (kg/m<sup>2</sup>) | 26.8 ± 4.3 | 26.8 ± 4.1 | 23.8 ± 4.0 |
| Triglycerides (mmol/L) | 1.5 ± 0.7 | 1.5 ± 0.7 | 1.0 ± 1.7 |
| Total cholesterol (mmol/L) | 5.7 ± 0.9 | 5.7 ± 0.9 | 4.7 ± 0.9 |
| HDL cholesterol (mmol/L) | 1.3 ± 0.3 | 1.3 ± 0.4 | 1.5 ± 0.4 |
| Friedewald LDL cholesterol (mmol/L) | 3.8 ± 0.8 | 3.8 ± 0.8 | 2.7 ± 0.8 |

Values are mean±SD.
\* Pooled results of eight cohorts from different geographical and ethnic backgrounds and age distributions; characteristics of each cohort are detailed in **Table S1**.

### Lipid and metabolite quantification

High-throughput NMR metabolomics was used to quantify 228 lipoprotein lipids and polar metabolite measures from serum or plasma samples in the PROSPER trial and eight cohorts by the Nightingale platform (Nightingale Health Ltd, Helsinki, Finland). This provides simultaneous quantification of routine lipids, particle concentration and lipid composition of 14 lipoprotein subclasses, abundant fatty acids, amino acids, ketones and glycolysis related metabolites in absolute concentration units (**Supplementary Table 2**) (31). The Nightingale NMR metabolomics platform has been widely used in epidemiological studies (32, 33) and the measurement methods has been described previously (31, 34, 35).

### Statistical analyses

The effects of statin therapy on the 228 metabolic measures in the PROSPER trial were assessed by comparing the mean metabolite concentrations in the treatment group to the placebo group 6 months after randomization. The between-group difference in concentration of each metabolic measure was quantified separately using linear regression with metabolite concentration as outcome and treatment status as predictor, adjusted for age and sex. All metabolite concentrations differences were scaled to standard deviation (SD) units to enable comparison of measures with different units and across wide ranges of concentration levels. Results in absolute units are presented in **Supplementary Table 3**. The percentage difference in metabolite concentration, relative to the placebo group, were examined as secondary analyses.

The effect of genetic inhibition of PCSK9 on each of the 228 metabolic measures was analyzed separately by fitting linear regression models with metabolite concentrations as outcome and rs11591147-T allele count as explanatory variable, representing the number of LDL-C lowering alleles. For sensitivity analysis, we conducted equivalent tests of each metabolic measure with rs12916-T in *HMGCR* as explanatory variable. All genetic analyses assumed an additive effect and were adjusted for age, sex, and the first four genomic principal components. Effect sizes and standard errors from each cohort were combined using inverse variance-weighted fixed effect meta-analysis. All effect sizes were scaled to SD units of metabolite concentrations, as for analyses of PROSPER. The similarity between the overall patterns of metabolic effects due to PCSK9 inhibition and statin therapy was summarized using the linear fit of the effect estimates of 153 metabolic measures (12), covering all measures except lipoprotein lipid ratios and five polar metabolites that could not be reliably quantified in PROSPER.

To facilitate comparison between the substantial metabolic effects of statin therapy with the smaller effects from genetic inhibition of PCSK9, results are presented relative to the respective lowering in LDL-C within each study design (in SD-units, as quantified by NMR metabolomics) (12, 35). For the statin trial, the estimates derived from comparing statin treatment to placebo were divided by 1.19; for PCSK9 genetic associations, per-allele effect estimates were divided by 0.44; for sensitivity analyses using rs12916 in *HMGCR*, per-allele effect estimates were divided by 0.078. The scaling relative to LDL-C allows us to interpret the reported effect sizes as a change in concentration in each metabolic measure (in SD units) that accompanies a 1-SD lowering of LDL-C by statin therapy and PCSK9 inhibitors.

Although 228 metabolic measures in total were examined, the number of independent tests performed is lower due to the correlated nature of the measures (35). The number of independent tests was estimated by taking the average number of principal components explaining 99% of the variation in the metabolic measures (35). Thus, significance was considered at P<0.0003 to account for the testing of 54 independent metabolic measures and three sets of analyses conducted (main effects of statins, *PCSK9*, and difference in their effects). To facilitate visualization of the results, we focused on 148 measures that cover all the metabolic pathways assayed; results for the remaining measures are provided in **Supplementary Figure 1** and **Supplementary Tables 4-6**. Statistical analyses were conducted using R3.2 (www.r-project.org).

## RESULTS

**Figure 1** provides an overview of the study design. Characteristics of the study populations are shown in **Table 1**. Characteristics of each of the eight cohorts used for the genetic analyses (N=72,185) are detailed separately in **Supplementary Table 1**. The detailed metabolic effects of statin treatment in PROSPER (pravastatin 40 mg/daily) and genetic inhibition of PCSK9 are compared in **Figure 2**. Overall, there was a high concordance of association of statin treatment and genetic inhibition of PCSK9 across the detailed metabolic profile (*R*^2^=0.88). Nonetheless, some discrepancies in effect sizes between statin treatment and *PCSK9* rs11591147 were evident, primarily for very-low-density lipoprotein (VLDL) lipids.

**Figure 2.**
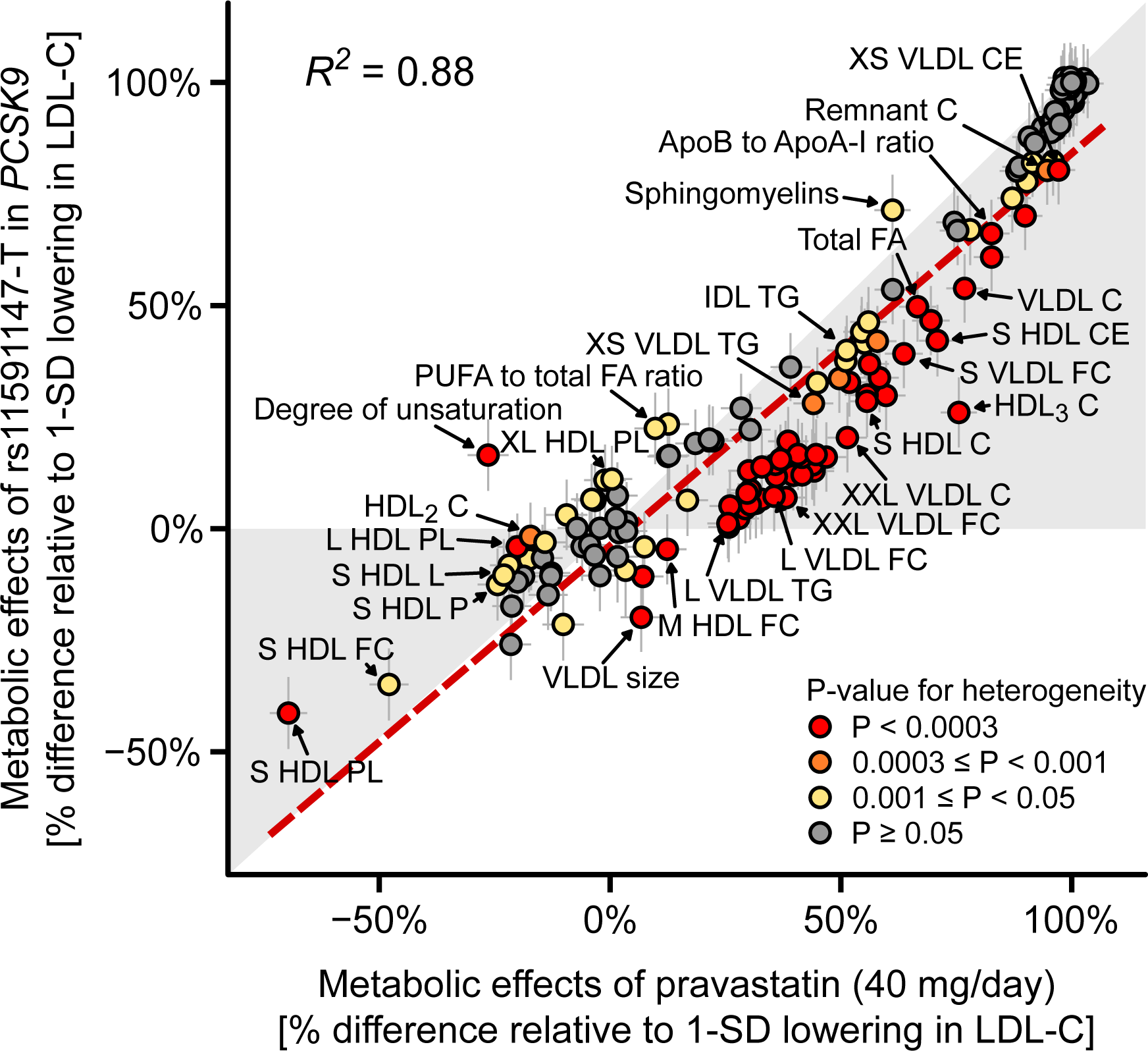
Consistency of metabolic effects of statin treatment and *PCSK9* rs11591147. Effect sizes of each metabolic measure is given with 95% confidence intervals in gray vertical and horizontal error bars. Color coding for the metabolic measure indicates the P-value for heterogeneity between statin therapy and *PCSK9* rs11591147. *R*^2^ indicates goodness of fit. The red dashed line denotes the linear fit between the metabolic effects (slope = 0.88). C: cholesterol; FA: fatty acids; HDL: high-density lipoprotein; IDL: intermediate-density lipoprotein; LDL: low-density lipoprotein; PL: phospholipids; PUFA: polyunsaturated fatty acids; TG: triglycerides; very-low-density lipoprotein. A full list of metabolite names are given in **Supplementary Table 2**.

### Effects on lipoprotein lipids

The specific effects of statin therapy and genetic inhibition of PCSK9 on lipid fractions and 14 lipoprotein subclasses are shown in **Figure 3**. Scaled to the same lowering of LDL-C, *PCSK9* rs11591147 displayed similar effects as statin therapy for total cholesterol and intermediate-density lipoprotein (IDL)-cholesterol, with no effect on high-density lipoprotein (HDL)-cholesterol. However, *PCSK9* rs11591147 had a weaker effect on lowering VLDL-cholesterol compared with statins (54% vs. 77%, relative to the lowering effect on LDL-C (%_LDL-C_); P_het_=2x10^-7^). These results were substantiated by the pattern of reduction in lipoprotein subclass particles: while the effects were similar for lowering particle concentrations in all three LDL subclasses, the extent of lowering of small, medium-sized and large VLDL particle concentrations was smaller for *PCSK9* rs11591147 compared with statin therapy. A similar discrepancy was observed for cholesterol concentrations within the six VLDL subclasses (**Figure 4**).

**Figure 3.**
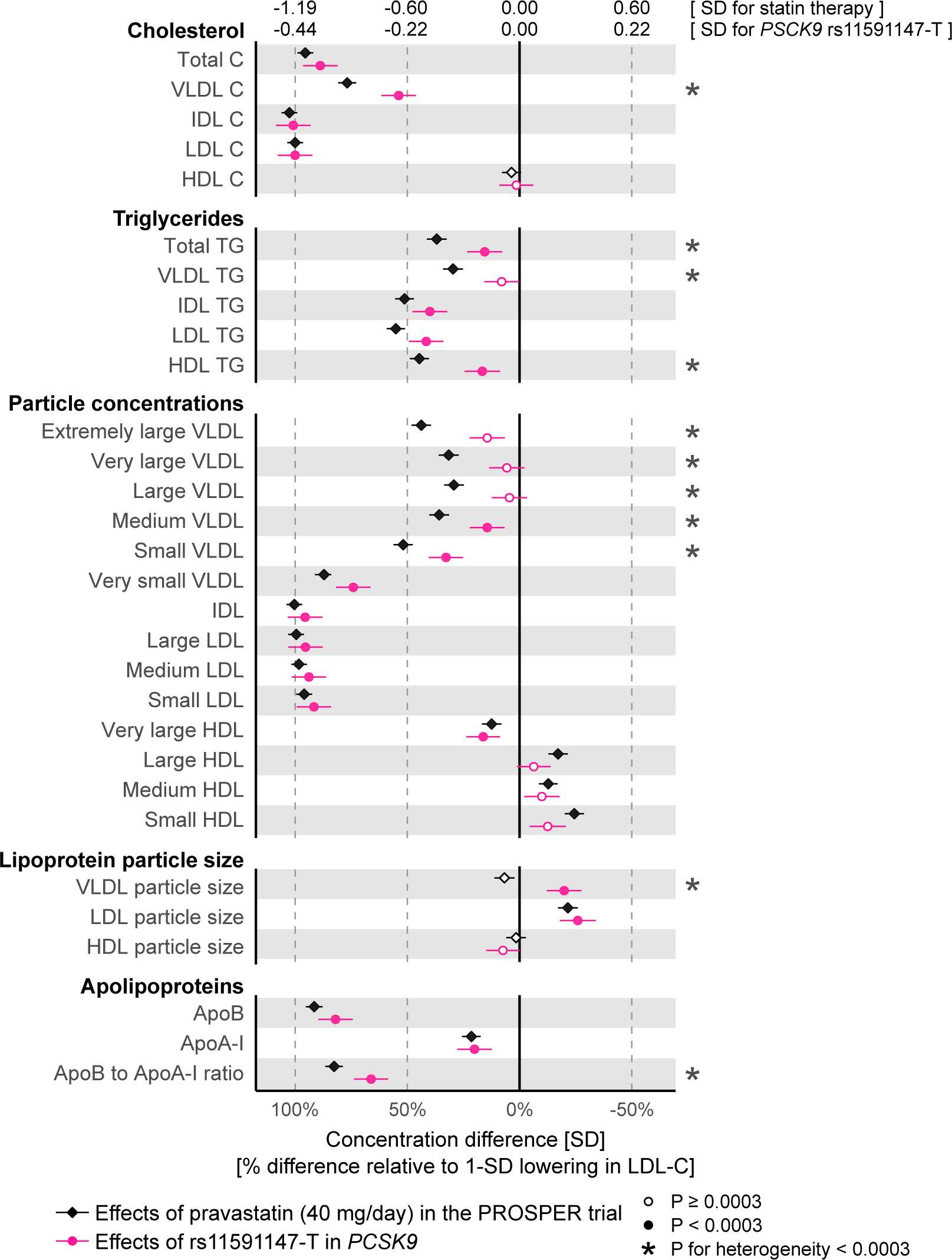
Effects of statin treatment and genetic inhibition of PCSK9 on lipoprotein and lipid levels. Differences in lipoprotein and lipid levels due to statin treatment were assessed in the PROSPER trial at 6-month post randomization (black diamonds; n=5359 for which 2659 were on pravastatin 40mg/day). The corresponding effects of *PCSK9* rs11591147 were assessed for n=72,185 by meta-analysis of eight cohorts (red circles). Error bars indicate 95% confidence intervals. Effect estimates are shown in SD-scaled concentration units (top axis) and relative to the lowering effect on LDL-C (bottom axis). The results for different lipid types within the 14 lipoprotein subclasses are shown in **Supplementary Figure 1**. Effects in absolute concentration units are listed in **Supplementary Table 3**.

**Figure 4.**
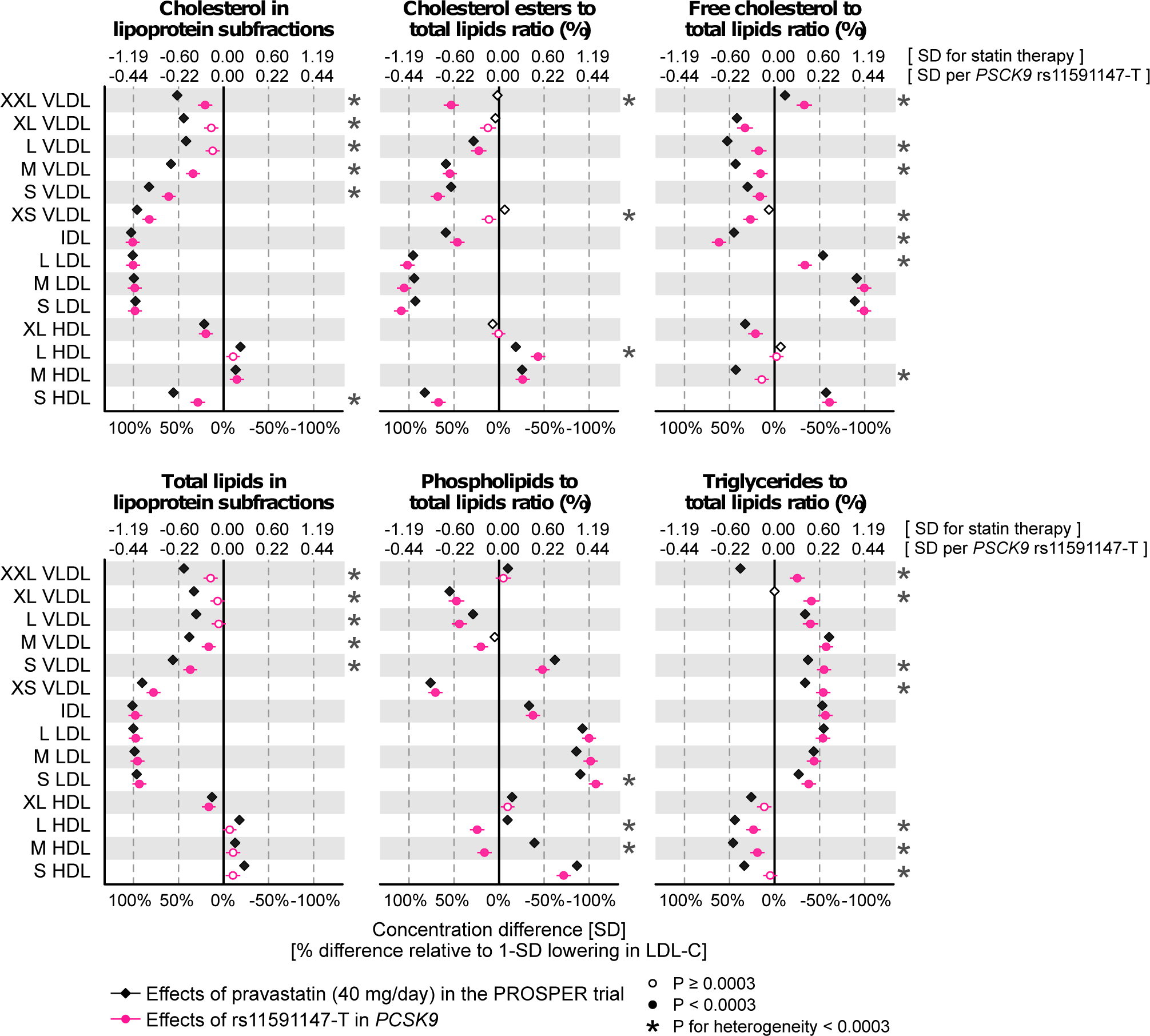
Effects of statin treatment and genetic inhibition of PCSK9 on lipoprotein composition. Differences in lipoprotein composition measures due to statin treatment were assessed 6-month post randomization in the PROSPER trial (black). The corresponding effects of *PCSK9* rs11591147 were assessed for n=72,185 (red). Error bars indicate 95% confidence intervals. Results are shown in SD-scaled concentration units (top axis) and relative to the lowering effect on LDL-C (bottom axis).

Both for statin therapy and genetic inhibition of PCSK9, the effects on triglyceride measures were modest compared to those observed for cholesterol levels in the same lipoprotein subfractions (**Figure 3**). Most pronounced lowering of triglycerides was seen for IDL and LDL particles. *PCSK9* rs11591147 displayed a weaker effect than statin therapy on lowering total plasma triglycerides (16%_LDL-C_ vs. 37%_LDL-C_; P_het_=3x10^-6^). Similar differences were seen for VLDL and HDL triglycerides. Consistent with the observed discrepancies for lowering of medium and large VLDL particles, genetic inhibition of PCSK9 resulted in modestly larger VLDL size whereas statin therapy had no effect on this measure. The effects on apolipoprotein concentrations were broadly similar, albeit a larger decrease was observed in the case of statins for the ratio of apolipoprotein B to A-I.

### Effects on lipoprotein composition

In addition to affecting the absolute lipid concentrations, both statin therapy and genetic inhibition of PCSK9 had prominent effects on the relative abundance of lipid types (free and esterified cholesterol, triglycerides, and phospholipids) in differently sized lipoprotein subclasses (**Figure 4**). The most pronounced lipoprotein composition effects were observed within LDL subclasses, with substantial lowering in the relative abundance of cholesteryl esters in LDL particles, alongside increases in the abundance of free cholesterol and phospholipids. These effects were very similar for statin treatment compared with *PCSK9* rs11591147. Subtle discrepancies between statin and genetic inhibition of PCSK9 were observed, e.g., for the extent of lowering the fraction of free cholesterol in VLDL particles. The relative fraction of triglycerides in LDL and other apolipoprotein-B carrying particles increased similarly for both statins and *PCSK9* rs11591147, whereas statin therapy caused larger decreases in the relative abundance of triglycerides within HDL.

### Effects on fatty acids and polar metabolites

The effects of statin therapy and genetic inhibition of PCSK9 on fatty acid concentrations and the balance of fatty acid ratios are shown in **Figure 5**. Absolute concentrations of all fatty acids were lowered, with the most pronounced lowering for concentrations of linoleic acid, an omega-6 fatty acid commonly bound to cholesteryl esters in LDL particles. The effects of statins and *PCSK9* rs11591147 were broadly similar, albeit with the lowering of total fatty acids being stronger in the case of statins (50%_LDL-C_ vs. 67%_LDL-C_; P_het_=2x10^-4^). The effects on the fatty acid ratios were generally modest, both for statin therapy and *PCSK9* rs11591147. A pronounced discrepancy between these was observed for the overall degree of fatty acid unsaturation (16%_LDL-C_ reduction for *PCSK9* vs. 26%_LDL-C_ increase for statin; P_het_=4x10^-20^).

**Figure 5.**
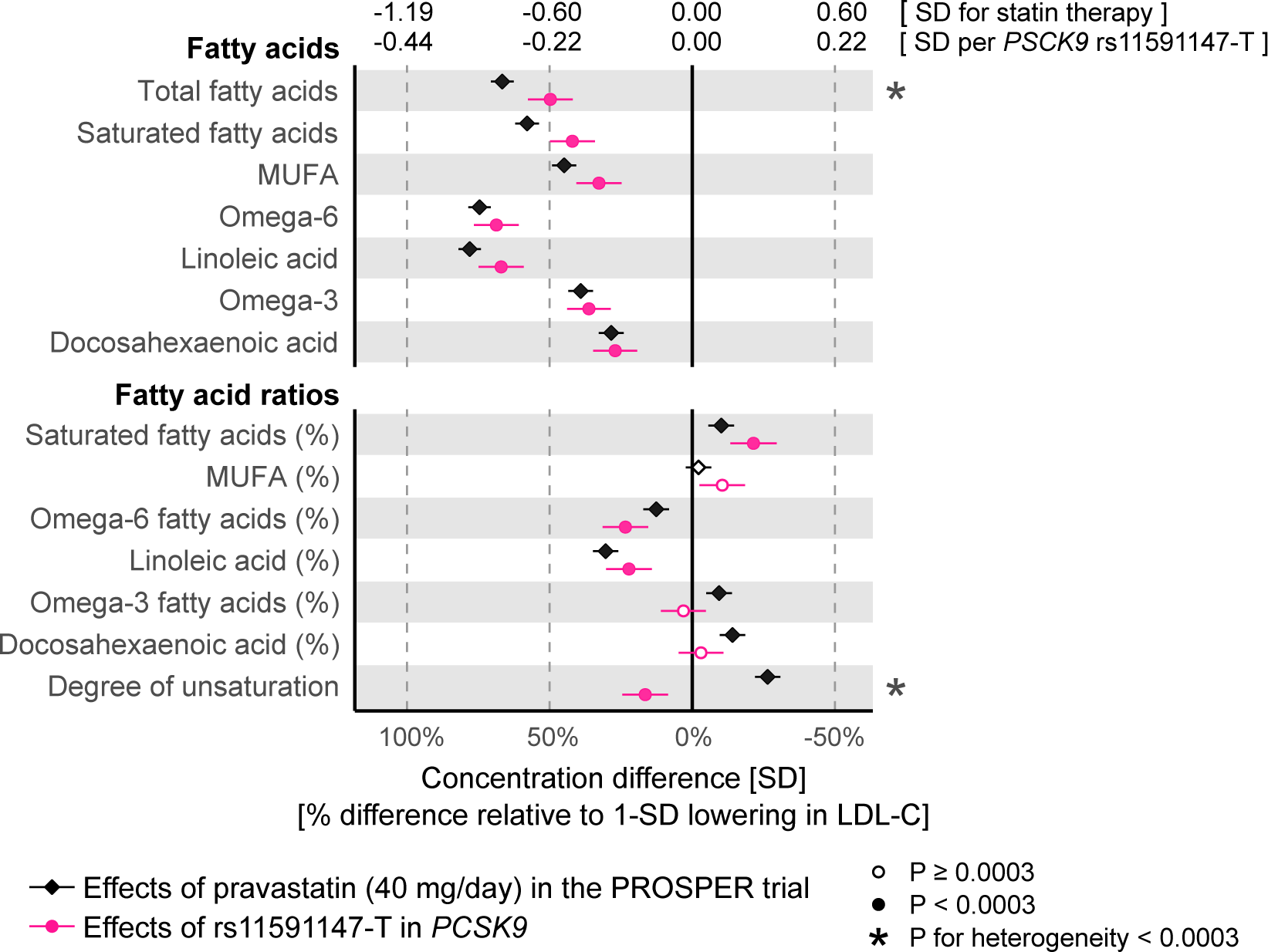
Effects of statin treatment and genetic inhibition of PCSK9 on fatty acids. Differences in fatty acid levels due to statin treatment were assessed 6-month post randomization in the PROSPER trial (black). The corresponding effects of *PCSK9* rs11591147 were assessed for n=72,185. Error bars indicate 95% confidence intervals. Results are shown in SD-scaled concentration units (top axis) and relative to the lowering effect on LDL-C (bottom axis).

We further assessed the effects of statin therapy and *PCSK9* rs11591147 on polar metabolites and other metabolic measures quantified simultaneously in the metabolomics assay, including circulating amino acids, glycolysis metabolites, ketone bodies, and GlycA, a marker of chronic inflammation (36) (**Figure 6**). Statin therapy caused only minor effects on these metabolic measures; the strongest lowering effects were observed for GlycA (17%_LDL-C_) and isoleucine (7%_LDL-C_). The effects of *PCSK9* rs11591147 were also very close to null for these measures, including for glycolysis related metabolites and markers of insulin resistance. Of note, information on glucose, lactate, and pyruvate were not available in the PROSPER trial due to glycolysis progression post-sample collection.

**Figure 6.**
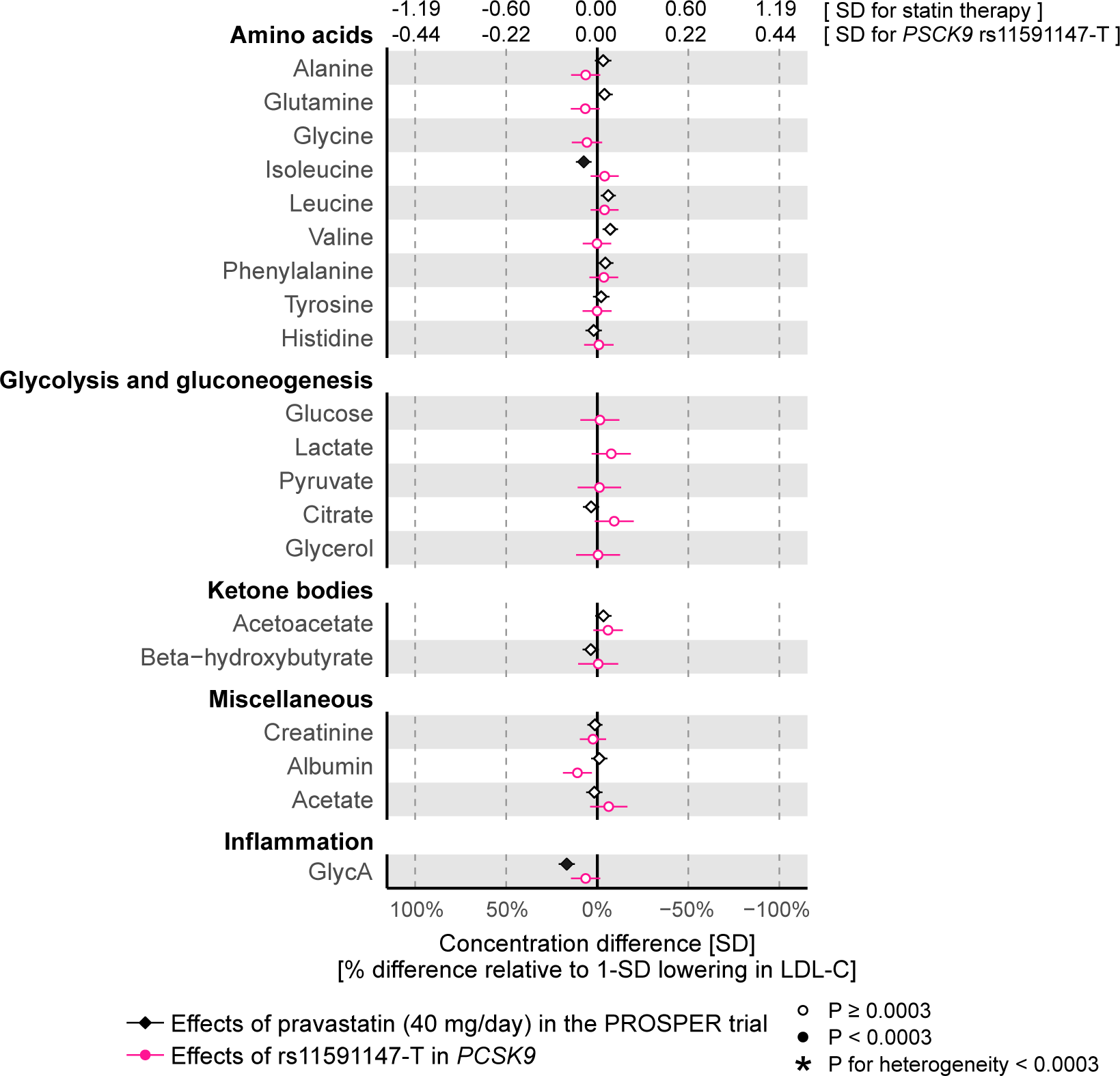
Effects of statin treatment and genetic inhibition of PCSK9 on polar metabolites. Differences in metabolite levels due to statin treatment were assessed 6-month post randomization in the PROSPER trial (black). The corresponding effects of *PCSK9* rs11591147 were assessed for n=72,185. Error bars indicate 95% confidence intervals. Glycine, glucose, lactate, pyruvate and glycerol measures were not available from PROSPER. Results are shown in SD-scaled concentration units (top axis) and relative to the lowering effect on LDL-C (bottom axis).

### Comparison to PREVEND-IT trial and Mendelian randomization

To replicate the detailed metabolic effects of statins observed in PROSPER, we compared them with recent results from the PREVEND-IT trial obtained using the same NMR metabolomics platform (13). PREVEND-IT also examined the effects of pravastatin (40mg/day), with metabolomic changes assessed from baseline to 3-month for 195 individuals on treatment. The detailed metabolic effects of statin treatment were highly concordant between PROSPER and PREVEND-IT (*R*^2^=0.96; **Supplementary Figure 2**). When results were scaled to an equivalent lowering in LDL-C, all significant discrepancies between effects of *PCSK9* rs11591147 compared to PROSPER were similar or somewhat larger in PREVEND-IT, with the exception of three measures of lipoprotein composition (**Supplementary Figure 1)**.

We further compared the metabolic effects of *PCSK9* rs11591147 to those caused by rs12916 in the *HMGCR* gene, hereby using the genetic variants to effectively act as two naturally occurring trials in the same study population. The overall pattern of metabolic effects was highly similar for *PCSK9* rs11591147 and *HMGCR* rs12916 (*R*^2^=0.92; **Supplementary Figure 3A**). Nonetheless, similar deviations in VLDL lipid measures were observed as when comparing *PCSK9* rs11591147 to the statin trial results (**Supplementary Figure 4**). Specifically, the lowering of particle concentrations in all VLDL subclasses, except very small VLDL, was more similar to statin treatment effects for *HMGCR* than for *PCSK9*. Similar subtle differences were observed for cholesterol and triglyceride concentrations in VLDL subclasses, whereas the lowering of total and saturated fatty acids were similar for the *HMGCR* and *PCSK9* variants. However, power to detect statistical differences on individual measures was modest due to the much weaker LDL-C lowering effect of *HMGCR* rs12916. Finally, the overall pattern of metabolic effects of statin therapy in PROSPER was highly concordant to effects of *HMGCR* rs12916 (*R*^2^=0.95; **Supplementary Figure 3B**), signifying pharmacological and genetic inhibition of HMG-CoA reductase, respectively.

In sensitivity analyses, the pattern of metabolic effects from *PCSK9* rs11591147 were similar across the eight cohorts (**Supplementary Figure 5**). We also observed similar detailed patterns of metabolic effects as for rs11591147 when examining other genetic variants in *PCSK9* that have previously been used in Mendelian randomization studies (11, 29)(**Supplementary Figure 6**). Results for all the 228 metabolic measures quantified are illustrated in **Supplementary Figure 1 and 4**. Metabolic effects in absolute concentration units are listed in **Supplementary Table 3**. The percentage differences in lipid and metabolite concentrations in the PROSPER statin trial are shown in **Supplementary Figure 7.** Exact effect estimates for all analyses are tabulated in **Supplementary Tables 4-6**.

## DISCUSSION

This study elucidates the comprehensive metabolic effects accompanying LDL-C reduction due to statin therapy and PCSK9 inhibition. The results demonstrate that, as compared to statin therapy, genetic inhibition of PCSK9 gives rise to comparable changes across many different markers of lipid metabolism. However, our results also suggest that, for a given LDL-C reduction, PCSK9 inhibitors may be somewhat less efficacious at lowering VLDL particles. This could potentially contribute to subtle differences in potency for cardiovascular event lowering for the same LDL-C lowering (6), since recent evidence suggests that VLDL-cholesterol and other triglyceride-rich lipoprotein measures may causally contribute to the development of coronary heart disease independent of LDL-C (37–39). Moreover, trial data show that VLDL-cholesterol is a stronger predictor of cardiovascular event risk than is LDL-C among patients on statin therapy (40, 41).

Statins and PCSK9 inhibitors both lower circulating LDL-C levels via upregulation of LDL receptors on cell surfaces. Consistent with this shared mechanism for clearance of LDL particles, we found that statins and genetic inhibition of PCSK9 caused a highly consistent pattern of change across the detailed metabolic profile. The metabolomic profiling of the PROSPER trial corroborates prior studies on the fine-grained metabolic characterization of statin effects, with exceptional consistency of results to those observed in longitudinal cohorts and a small randomized trial (12, 13). By metabolomic profiling a large number of individuals from multiple cohorts, our results also validate and extend previous studies examining the detailed metabolic effects of *PCSK9* rs11591147 (14, 15). Importantly in relation to assessment of potential side-effects of PCSK9 inhibition, we did not observe effects on amino acids or other non-lipid metabolites, despite many of these biomarkers showing associations with risk of incident diabetes and cardiovascular risk (32, 42, 43). Our results on the pattern of lowering VLDL particles due to genetic inhibition of PCSK9 are in agreement with two small trials assessing the effects of the PCSK9 inhibitors alirocumab and evolocumab on lipoprotein particle concentrations; both trials showed substantial reduction in small and medium-sized VLDL particles, while the particle concentration of large VLDL fraction was not affected (16, 17). Similar results were also found in a small PCSK9-inhibitor trial using separation of VLDL subfractions and other lipid measures by ultracentrifugation, which also corroborate our results on a stronger effect on lowering of VLDL-cholesterol as compared to total plasma triglycerides (18). However, differences in assay methods complicate direct comparison of these trials to our results. In combination, these results provide orthogonal evidence for diverse lipoprotein lipid alterations by PCSK9 inhibitors, coherent with the comprehensive metabolic effects of statins.

Currently licensed PCSK9 inhibitors are given either instead of statins – when there is strong evidence of statin intolerance in those with familial hypercholesterolemia – or on top of maximally tolerated statins including in patients with existing vascular disease (44). Such treatment with PCSK9 inhibitors has been shown to be more efficacious in lowering LDL-C than the most potent statins (4–6, 8). Mendelian randomization studies comparing *PCSK9* and *HMGCR* gene scores on cardiovascular outcomes have indicated nearly identical protective effects per same LDL-C lowering (11, 45). However, when scaling the metabolic effects to an equivalent lowering in LDL-C, our results indicate subtle differences in multiple lipoprotein lipid measures. The most notable discrepancy was for VLDL lipids, suggesting weaker potency of PCSK9 inhibitors in clearance of these triglyceride-rich lipoproteins as compared with statins. The causal consequences of these differences in medium-sized and large VLDL particles, that are rich in triglycerides, remains unclear and warrants further investigations; whereas IDL and the smallest VLDL particles can penetrate the arterial wall to cause atherosclerosis, it is commonly perceived not be to the case for larger VLDL particles (37, 46). We also observed a difference in lowering of VLDL-cholesterol levels; the cholesterol concentrations of VLDL particles are strongly associated with risk of myocardial infarction (43) and some studies have suggested that VLDL-cholesterol could underpin the link between triglycerides and cardiovascular risk (37, 40). If these VLDL particles do play a causal role in vascular disease, the discrepancy between statin therapy and PCSK9 inhibition could translate into slightly more potent cardiovascular risk reduction for the same LDL-C lowering for statins as compared with PCSK9 inhibition. We acknowledge that the present comparison of detailed metabolic effects of statin therapy and PCSK9 inhibition does not inform on the cardiovascular benefits of anti-PCSK9 therapies above current optimal care, but potentially in keeping with our findings, the first major cardiovascular outcome trial on PCSK9 inhibition did demonstrate slightly weaker cardiovascular event lowering compared to meta-analysis of statin trials per mmol/L reduction in LDL-C (6). While likely explanations for this discrepancy include the short trial duration (8) and choice of primary end-point, other explanations, such as differences in anti-inflammatory effects, have also been suggested (11, 47). Our results provide an additional hypothesis for exploration: the apparent weaker cardioprotective effects of PCSK9 inhibitors compared to statins per unit reduction in LDL-C may be due to weaker reductions in VLDL lipid concentrations by PCSK9 inhibition. This hypothesis warrants further investigation, including elucidation of the causal role of triglyceride-rich VLDL particles in tandem with further examinations of the detailed lipid effect of PCSK9 inhibitors.

Strengths and limitations of the study should be considered. The lack of NMR metabolomics data for a PCSK9 inhibition trial motivated the use of a loss-of-function variant in *PCSK9*, as a proxy for the anticipated therapeutic effects. The close match in the detailed metabolic effects of statin therapy and *HMGCR* observed in this study, substantiates the validity of using genetic variants to mimic lipid-lowering effects in randomized trial settings. While we note that the metabolic profile of other statins may differ to that of pravastatin, the similarity between *HMGCR* and statin therapy we identified provides reassurances to the generalizability of our findings. To robustly assess the metabolic effects of genetic inhibition of PCSK9, we had more than five times the sample size of prior studies examining *PCSK9* rs11591147 on detailed lipoprotein subclass profiles (14, 15). Despite the large sample size, we had limited power to detect effects on glycolysis-related metabolites due to pre-analytical effects causing depletion of glucose levels in the blood samples. A strength of the metabolomics platform used is the ability to profile lipoprotein subclasses and their lipid composition at high-throughput, however we acknowledge that other assays may provide even deeper characterization of lipid metabolism and non-lipid pathways to further clarify the molecular effects of lipid-lowering therapies (48).

In conclusion, we found highly similar metabolic effects of statin therapy and genetic inhibition of PCSK9 across a comprehensive profile of lipids, lipoprotein subclasses, fatty acids, and polar metabolites. The detailed profiling of lipoprotein subclasses revealed weaker effects of PCSK9 inhibition on VLDL particles and their cholesterol concentrations as compared with statins, when scaled to an equivalent lowering of LDL-C. If some of these VLDL lipids have independent causal effects on cardiovascular risk, this could contribute to subtle differences in cardiovascular event reduction between statins and PCSK9 inhibitors. More broadly, these results highlight the need for large-scale metabolomics in combination with randomized trials and genetics to uncover potential molecular differences between related therapeutics.

## DISCLOSURES

PW is employee and shareholder of Nightingale Health Ltd, a company offering NMR-based metabolic profiling. JK reports stock options in Nightingale Health. DAL has received support from several government and charity health research funders and from Roche Diagnostics and Medtronic for research unrelated to that published here. The Clinical Trial Service Unit & Epidemiological Studies Unit (MVH, RW, KL, IM, RC, ZC) has received research grants from Abbott/ Solvay/Mylan, AstraZeneca, Bayer, GlaxoSmithKline, Merck, Novartis, Pfizer, Roche, and Schering. MVH has collaborated with Boehringer Ingelheim in research, and in accordance with the policy of the The Clinical Trial Service Unit and Epidemiological Studies Unit (University of Oxford), did not accept any personal payment. NS has consulted for AstraZeneca, Bristol-Myers Squibb, Amgen, Sanofi, and Boehringer Ingelheim. No other authors reported disclosures.

## SOURCES OF FUNDING

This study was supported by the Academy of Finland (grant numbers 312476, 312477, 297338 and 307247), University of Oulu Graduate School, Strategic Research Funding from the University of Oulu, Finland, the Novo Nordisk Foundation (grant number NNF17OC0026062 and 15998), the Sigrid Juselius Foundation, and the UK Medical Research Council via the MRC University of Bristol Integrative Epidemiology Unit (MC_UU_12013/1 and MC_UU_12013/5).

PROSPER metabolic profiling by NMR was supported by the European Federation of Pharmaceutical Industries Associations (EFPIA), Innovative Medicines Initiative Joint Undertaking, European Medical Information Framework (EMIF) grant number 115372 and the European Commission under the Health Cooperation Work Programme of the 7th Framework Programme (Grant number 305507) “Heart ‘omics’ in AGEing” (HOMAGE).

The INTERVAL academic coordinating centre receives core support from the UK Medical Research Council (G0800270), the BHF (SP/09/002), the NIHR, and Cambridge Biomedical Research Centre, as well as grants from the European Research Council (268834), the European Commission Framework Programme 7 (HEALTH-F2-2012-279233), Merck, and Pfizer. The INTERVAL study is funded by NHSBT (11-01-GEN) and has been supported by the NIHR-BTRU in Donor Health and Genomics (NIHR BTRU-2014-10024) at the University of Cambridge in partnership with NHSBT.

ALSPAC receives core support from the UK Medical Research Council and Wellcome Trust (Grant: 102215/2/13/2) and the University of Bristol. Data collection and metabolic profiling in the ALSPAC mother’s study were obtained from British Heart Foundation (SP/07/008/24066) and the Wellcome Trust (WT092830M). Genetic data in the ALSPAC mothers was obtained through funding from the Wellcome Trust (WT088806). ALSPAC offspring genetic data was obtained with support from 23andMe.

The China Kadoorie Biobank baseline survey and first re-survey was supported by the Kadoorie Charitable Foundation in Hong Kong. Long-term follow-up has been supported by the UK Wellcome Trust (202922/Z/16/Z, 088158/Z/09/Z, 104085/Z/14/Z), National Key Research and Development Program of China (2016YFC0900500, 2016YFC0900501, 2016YFC0900504), Chinese Ministry of Science and Technology (2011BAI09B01), and National Natural Science Foundation of China (Grants No. 81390540, No. 81390541, No. 81390544). NMR metabolomics was supported by the BHF Centre of Research Excellence, Oxford (RE/13/1/30181). The British Heart Foundation, UK Medical Research Council and Cancer Research UK provide core funding to the Clinical Trial Service Unit and Epidemiological Studies Unit (CTSU) at the University of Oxford. MVH was supported by the National Institute for Health Research (NIHR) Oxford Biomedical Research Centre. We thank the CKB participants and the project staff based at Oxford, Beijing, the 10 regional centres, and the China National CDC and its regional offices. DNA extraction and genotyping was performed by BGI, Shenzhen, China.

The FINRISK studies have received financial support related to the present study from the he National Institute for Health and Welfare, the Academy of Finland (139635), and the Finnish Foundation for Cardiovascular Research (to V.S).

The Northern Finland Birth Cohorts of 1966 and 1986 has received financial support from Academy of Finland, University Hospital Oulu, Biocenter Oulu, University of Oulu, the European Commission (Horizon2020: 633595), the UK Medical Research Council, and Wellcome Trust.

The funders of the individual study cohorts had no role in the design and conduct of the study; collection, management, analysis, and interpretation of the data; preparation, review, or approval of the manuscript; and decision to submit the manuscript for publication. The views expressed in this paper are those of the authors and not necessarily any funding body.

## SUPPLEMENTARY MATERIAL

Supplementary Methods

Table S1. Clinical characteristics of the eight cohorts included in the genetic analyses of *PCSK9* rs11591147 and *HMGCR* rs12916

Table S2. Metabolite mean (SD) concentrations in PROSPER and cohorts used for genetic analyses

Figure S1. Effects of statin treatment and genetic inhibition of PCSK9 on 228 metabolic traits.

Figure S2. Consistency of metabolic effects of pravastatin 40 mg/day in PROSPER vs. PREVEND IT trials.

Figure S3. Consistency of metabolic effects of *HMGCR* rs12916-T vs. (A) *PCSK9* rs11591147-T, and (B) statin therapy in the PROSPER trial.

Figure S4. Metabolomic effects of statin treatment, *HMGCR* rs12916-T and *PCSK9* rs11591147-T.

Figure S5. Consistency of metabolomic effects of *PCSK9* rs11591147-T across the cohorts on 228 metabolic traits.

Figure S6. Consistency of metabolic effects of the main instrument *PCSK9* rs11591147-T with other *PCSK9* variants.

Figure S7. Percentage differences in metabolite concentrations in pravastatin versus placebo group in PROSPER trial.

### Spreadsheet file

Table S3. Main results in absolute concentration units.

Table S4. Metabolic effects of statin therapy in the PROSPER trial in SD-units.

Table S5. Metabolic effects of *PCSK9* rs11591147-T in SD units.

Table S6. Metabolic effects of *HMGCR* rs12916-T in SD-units.

